# Integrin cytoplasmic domain and pITAM compete for spleen tyrosine kinase binding

**DOI:** 10.1101/524447

**Authors:** Lina Antenucci, Maarit Hellman, Vesa P. Hytönen, Perttu Permi, Jari Ylänne

## Abstract

In hematopoietic tissues cell-cell communication involves immunoreceptors and specialized cell adhesion receptors that both mediate intracellular signals. Spleen tyrosine kinase (Syk) is a non-receptor tyrosine kinase involved in the downstream signaling of both immunoreceptors tyrosine activation motif (ITAM) receptors and integrin family cell adhesion receptors. Both phosphorylated ITAM (pITAM) and integrins bind to the regulatory domain of Syk composed of two Src homology 2 (SH2) domains. The interaction with pITAM is mediated by binding of a specific phosphotyrosine to each of the SH2 domains, leading to conformational changes and Syk kinase activation. Integrins bind to the interdomain A segment between the two SH2 domains and to the N-terminal SH2 domain, but the detailed binding site is not known. In order to map the binding site, we performed NMR titration experiments. We found that integrin cytoplasmic domain peptide induced chemical shift changes near the IA segment and at the phosphotyrosine binding site of the N-terminal SH2 domain of Syk. These changes were distinct, but partially overlapping with those induced by pITAM peptide. We were also able to show that pITAM peptide inhibited integrin binding to Syk regulatory domain. These results suggest that ITAM receptors and integrins cannot bind simultaneously to Syk, but provide two distinct routes for Syk activation.

---

Tyrosine kinases are an important class of cellular signaling enzymes and main targets of several cancer drugs (1). Understanding the regulation of tyrosine kinases is a key for understanding the function of cellular signaling networks. All non-receptor tyrosine kinases share a similar enzymatically active kinase domain but their regulatory domains allow their classification to 10 functional groups (2). Spleen tyrosine kinase (Syk) is a non-receptor tyrosine kinase that belongs to a distinct kinase subfamily whose regulatory domain is composed of two phosphotyrosine-binding Src homology 2 (SH2) domains (3). The other member of this subfamily in mammals is the zeta-chain associated kinase of 70 kD (ZAP70) (3). Gene depletion studies in mouse have shown that Syk is specifically required for the development of B-cells (4, 5) and ZAP70 for T-cell maturation (6, 7). Syk-deficient mice die newborn because of failure to separate blood vessels and lymphatic vessels (4, 5), a defect that may reflect the role of platelet adhesion in this process (3). In addition to platelets and lymphocytes, Syk has essential function in many other cells of hematopoietic origin as well as in many non-hematopoietic tissues, for instance in breast cancers (8). This reflects its function in mediating signal transduction from adaptive immune receptors, innate immune receptors and certain cell adhesion receptors.

The mechanism of Syk activation downstream of most of the above listed receptors is rather well understood: Src family kinases (Fyn and Lck) are activated upon immune receptor clustering and phosphorylate two Tyr residues in the cytoplasmic domains of the immune receptor complex (reviewed in (3)). These residues are found in a conserved sequence motif called the immunoreceptor tyrosine-containing activation motif (ITAM) that has a consensus sequence: YxxL/Ix6-12YxxL/I. ITAM is found either in the main transmembrane spanning polypeptide of the immune receptor (as in FcγRIIA) or in a separate transmembrane adaptor in immune receptors (as in TCR, BCR and FcεRI), innate immune pattern-recognizing receptors (in certain C-type lectin receptors as CLEC4E, CLEC6A, CLEC5A), or cell adhesion receptors (in platelet glycoprotein IV, GPIV, and osteoclasts-associated receptor, OSCAR). In addition, C-type lectin receptors exists where the full ITAM motif is formed by clustering of receptors containing single phosphorylated tyrosine motifs, so called hemITAM motifs (CLEC2, CLEC7A, CLEC9A) (reviewed in (3)). Phosphorylated ITAM (pITAM) sequences are able to interact with the two SH2 domains of Syk so that each of the pTyr residues binds to one SH2 domain (9). This interaction changes the relative orientation of these two domains resulting in the detachment of interdomain A (IA) and interdomain B (IB) segment from the kinase domain leading to kinase domain activation (9, 10).

In contract to phosphorylation-dependent interaction with transmembrane receptors or adaptors, described above, Syk interacts with integrin family cell adhesion receptors independently on receptor phosphorylation (11, 12). Direct interaction has been shown at least with β_1_, β_2_ and β_3_ integrin subunit cytoplasmic domains (12, 13). Integrin cytoplasmic domains contain Tyr residues, but they do not form ITAM motifs and Tyr phosphorylation of integrin tails inhibits Syk interaction (12, 13).

In many cell adhesion events ITAM signaling and integrin signaling happens simultaneously and sometimes even in the same adhesion complex. For instance, during platelet adhesion to vessel wall, ITAM-dependent signaling from the GPIV collagen receptor may happen simultaneously with the function of the integrin α_2_β_1_ collagen receptor (14) and is immediately followed by the function of integrin α_IIb_β_3_, which is the major fibrinogen receptor mediating platelet aggregation but also interact with several other extracellular matrix proteins (15). In immunological synapses TCR and α_L_β_2_ integrins function simultaneously in specific adhesion zones and signaling from these zones or cluster is regulated by diverse kinases and phosphatases (16). During osteoclast adhesion to bones, ITAM-linked OSCAR and integrin α_V_β_3_ signal parallelly (17).

While the pITAM-Syk interaction mechanism is well known, molecular details of integrin-Syk interaction are still unclear. Even though there are variations of relative affinities of various pITAM sequences binding to individual SH2 domains (18), present evidence shows that pITAM interaction with SH2 domains mostly follow the general mode of pTyr/SH2 interaction (9, 19). The main interaction is mediated by the phosphate group of the pTyr residue and in addition hydrophobic interaction with the +3 position Ile or Leu residue determines the specificity of the interaction (19). The exact residues of integrin tail mediating the contact with Syk regulatory domain are not known, but truncation studies have shown that the deletion of four C-terminal residues in β_3_ abolish the interaction (12) and that the minimum β_3_ cytoplasmic tail peptide is Arg734-Thr762 (29 residues) (12). On the Syk side, the main interaction sites have been mapped to the interdomain A (IA) segment and the N-terminal SH2 domain (13)

We have earlier shown that binding of soluble β_3_ integrin cytoplasmic domain does not directly activate Syk as in the case of soluble pITAM, but clustered integrin peptides can activate purified Syk (20). This fits well with the idea that integrins do not bind to the two SH2 domains in a similar way as pITAM and thus they cannot induce similar reorientation of Syk SH2 domains as pITAM.

In this study we set to test the hypothesis that integrin and pITAM binding to Syk are independent on each other. We used surface plasmon resonance (SPR) and nuclear magnetic resonance (NMR) spectroscopy methods to compare the binding surfaces of pITAM and β_3_ integrin cytoplasmic domain on Syk regulatory domains. We find that the Syk IA segment is the main binding site of integrin, but on the N-terminal SH2 domain, binding of integrin and pITAM peptides induce NMR chemical shift perturbations (CSPs) on overlapping areas. Interestingly, the integrin responsive surface on N-SH2 is close to the location of the IA in published structures. We also show that soluble pITAM inhibits Syk regulatory domain binding to integrin-coated surfaces. We believe these findings are important for understanding the cross-talk between integrins and ITAM receptors in Syk signaling.

## Results

### Mapping of pITAM and integrin binding surfaces on the N-terminal SH2 domain of Syk

To be able to study the possible cross-talk between integrin cytoplasmic domain and pITAM binding, we first set up to verify the binding site of integrin β_3_ peptide on Syk. We purified four fragments of the regulatory domain of Syk. The tandem SH2 (tSH2) fragment contained both SH2 domains and the intervening interdomain A (IA) segment. We also used either N-terminal SH2 (N-SH2) or the C-terminal SH2 (C-SH2) domains with the IA segment or N-SH2 alone (Figure 1). A 32-residue peptide from the C-terminus of integrin β3 subunit was coupled to the SPR sensor chip via an N-terminal Cys residue and the protein fragments were injected to the soluble phase. We found that the tSH2 and N-SH2+IA fragments bound integrin with similar dissociation constants (K_D_:s) of 3 μM and 8 μM, respectively (Figure 2A,B). The N-SH2 domain bound remarkably weaker (K_D_ = 130 μM) and IA-C-SH2 fragment somewhat weaker (K_D_=25 μM) to the integrin β_3_ peptide (Figure 2C, D). The K_D_, k_on_ and k_off_ values are shown in Table 1. We were not able to test the C-SH2 domain alone due to protein aggregation. The results are consistent with earlier observations (13) i.e. the IA segment of Syk is the major interaction site for β_3_ integrin tail and the N-SH2 domain contributes to the interaction.

**Figure 1:**
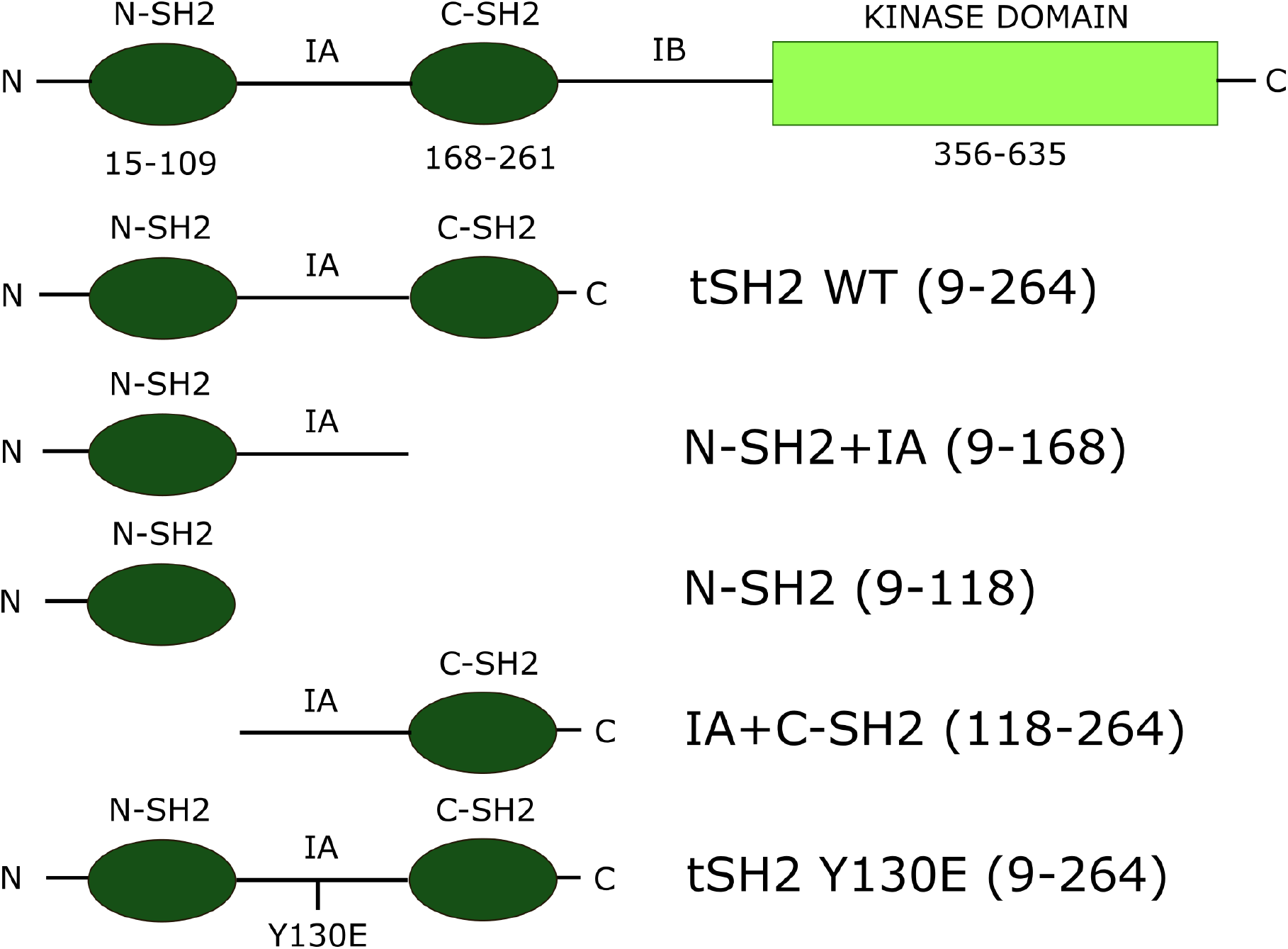
Schematic representation of tSH2 constructs used in this study. The SH2 domains are represented in dark green and the kinase domain in light green. The scheme of full length Syk is present as reference, but it was not used in the current study.

**Figure 2:**
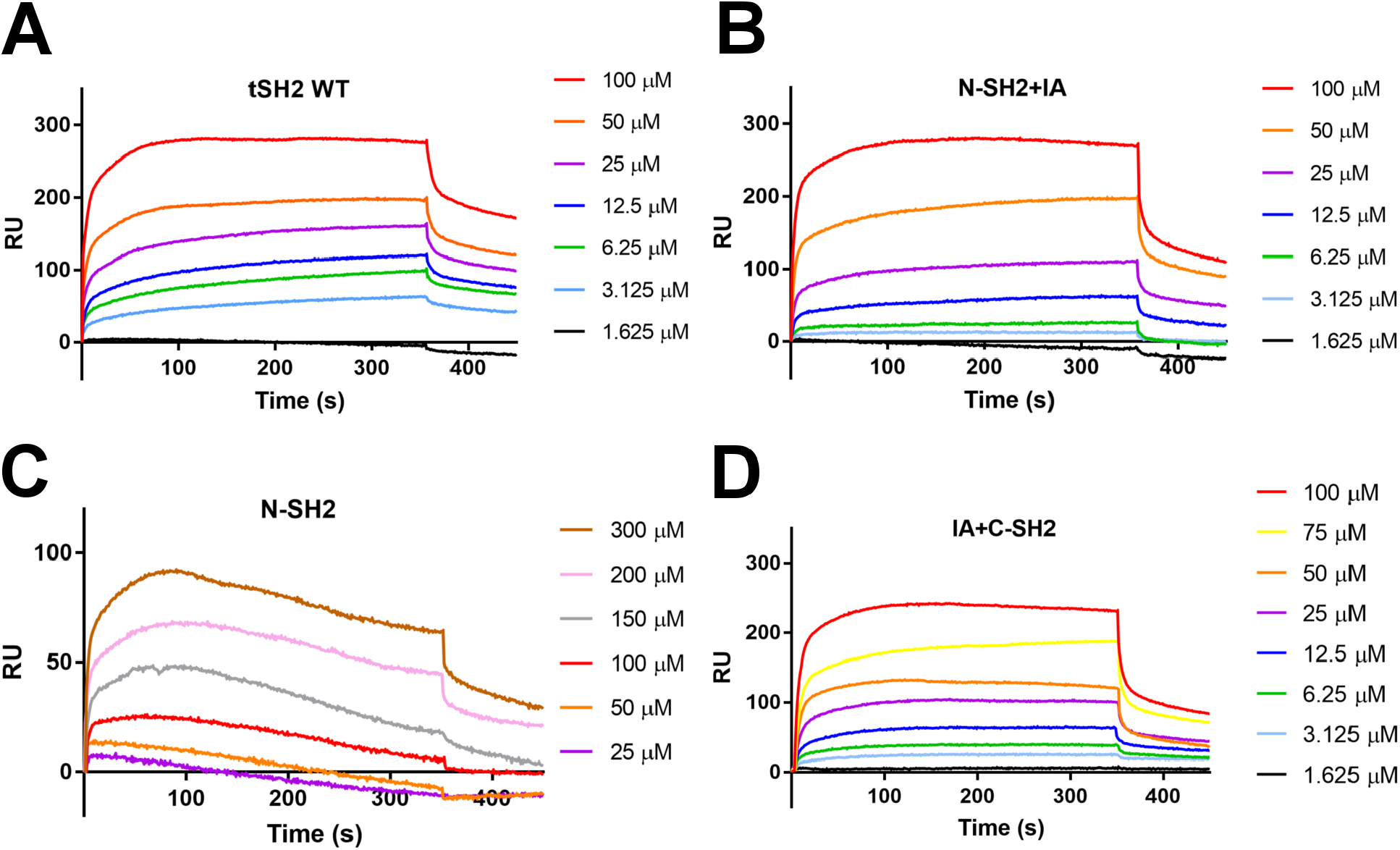
Interaction between integrin β3 cytoplasmic domain with tSH2 and shorter tSH2 constructs. Surface plasmon resonance experiments where integrin β3 cytoplasmic domain peptide was coupled on the surface (ligand) and different protein concentrations (from 1.625 to 300 μM) have been injected (analyte). In the graphs, for each construct, the variation of Response Units (RU) in the time (s) is plotted. The injection/association phase starts at time 0 seconds and ends in the time point marked with an *arrow;* the end of the injection coincides with the beginning of the dissociation phase. The graphs where fitted with Langmuir model using Bia Evaluation software to calculate the K_D_ and kinetic constants.

**Table 1:** Binding constants of tSH2 constructs calculated from the surface plasmon resonance experiments, where the biosensor surface was functionalized with integrin β3 cytoplasmic tail peptide

| | $k_{on}(1/M*s)$ | $k_{off}(1/s) * 10^{-3}$ | $K_D(M) * 10^{-6}$ |
| --- | --- | --- | --- |
| tSH2 wild type | 64.0±1.81 | 0.20±0.13 | 3.13±0.71 |
| N-SH2+IA | 60.5±1.76 | 0.50±0.11 | 8.33±0.83 |
| N-SH2 | 123±18.5 | 16.1±1.28 | 131±3.17 |
| IA+C-SH2 | 236±5.79 | 6.11±0.09 | 25.9±0.09 |

We further analyzed the binding site on pITAM and the β3 integrin peptide using solution state NMR spectroscopy. We carried out the chemical shift assignment of N-SH2 fragment (Antenucci et al., manuscript in preparation) and transferred the assignments to the ^15^N, ^1^H correlation spectrum (^15^N-HSQC) of N-SH2+IA. Hemi-phosphorylated pITAM peptide was titrated to the ^15^N-labeled N-SH2 fragment and the binding induced CSPs were analyzed (Figure 3A, B). The major changes mapped to the crystallographically determined binding site of pITAM (Figure 3C). To map the integrin binding surface on Syk, the ^15^N-labeled N-SH2+IA fragment was used instead, given that the integrin peptide was not well soluble in concentrations above 500 μM and the K_D_ for its binding to N-SH2 alone was estimated to be 130 μM. On the contrary, K_D_ with N-SH2+IA was 8 μM, as determined using SPR. NMR titration with the β_3_ integrin peptide revealed distinct CSPs for residues 20-23 and 44-51 (Figure 3E,F). In addition to the observed CSPs for the assigned peaks in the N-SH2 domain, some unassigned peaks were also shifted upon interaction (Supporting information figure). These unassigned peaks were apparently derived from the IA segment indicating that they are involved in binding of the β_3_ integrin peptide. Interestingly, some of the residues in the N-SH2 domain showed CSPs both in the presence of β3 integrin and pITAM. Particularly only Arg22 main chain NH resonance was shifted by integrin, whereas only its side chain N^ε^-H^ε^ was shifted by pITAM. This is in good accordance with the crystal structure of the tSH2-pITAM complex, where N^η^ of Arg22 interacts with phosphate group of pITAM (9). Arg45 main chain NH and Asn46 side chain NH2 were shifted by both integrin and pITAM. Visualization these shifts on the surface of the domain (Figure 3A,E) suggests that the interaction sites of pITAM and integrin overlap partially on the N-SH2 domain of Syk.

**Figure 3:**
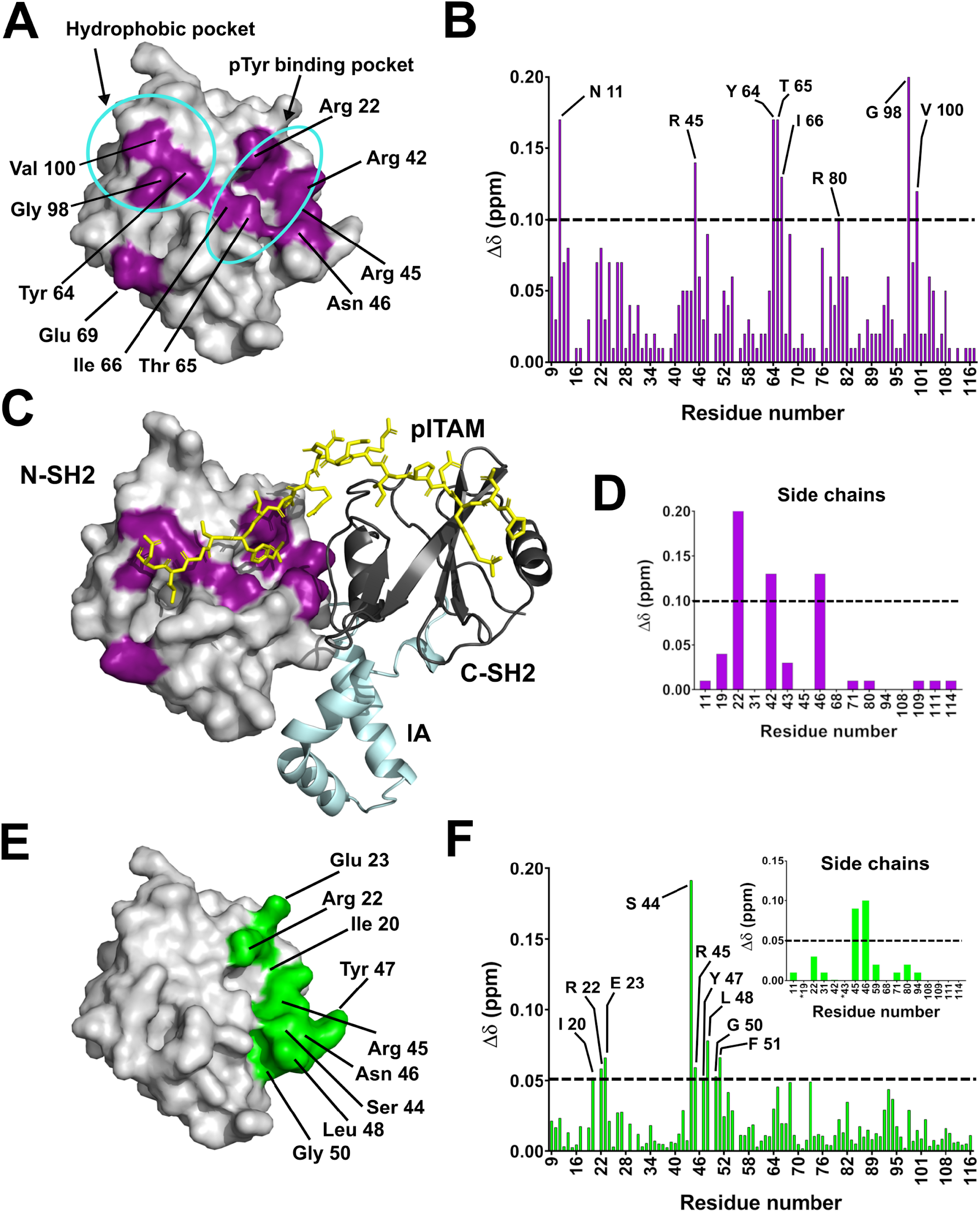
NMR chemical shift perturbations induced by ppITAM and integrin peptides. **A.** Surface representation of N-SH2 structure (PDB: 4FL2). The amino acids, whose chemical shifts changed upon addition of ppITAM peptide are highlighted in purple. **B.** Bar graph showing the chemical shift perturbations (Δδ, ppm) observed in ^15^N-HSQC spectra of the N-SH2 in presence of ppITAM (1:80, protein:peptide ratio). The shifts higher than 0.1 were considered significant and the cut off line is indicated. **C.** Comparison of the crystal structure of ppITAM-tSH2 complex (PDB: 1A81) with the chemical shift perturbations shown in A. **D.** Chemical shift perturbations in the ^15^N-HSQC spectra of the N-SH2 side chains in from the same titration as A. **E.** Amino acids showing a shift higher than 0.05 with the integrin peptide mapped on the same structure as in A and shown in green. **F.** Bar graph showing the chemical shift change (Δδ, ppm) observed in ^15^N-HSQC spectra of the N-SH2 backbone in presence of integrin peptide (protein:peptide ratio 1:20). The cut off line at 0.05 is indicated. The side chain shifts are shown in the insert.

### pITAM inhibits integrin binding to N-SH2 domain of Syk

To test if pITAM and integrin β_3_ cytoplasmic peptide compete for binding to the regulatory domain of Syk, we performed SPR competition experiments. pITAM was able to inhibit tSH2 binding to integrin-coated SPR surface with IC50 value of 15 μM (Figure 4A). To make sure that pITAM concentrations used in this experiment were high enough for saturation binding, we measured the K_D_ of pITAM to the same tSH2 preparation using thermofluor methods. We found that pITAM stabilized the melting temperature of tSH2 and this stabilization could be fitted to a single site binding equation with K_D_ of 4.2 ± 0.68 μM (Figure 4B,C). These data suggest that, contrary to our initial hypothesis, pITAM and integrin cytoplasmic domain compete for binding to the regulatory domain of Syk.

**Figure 4:**
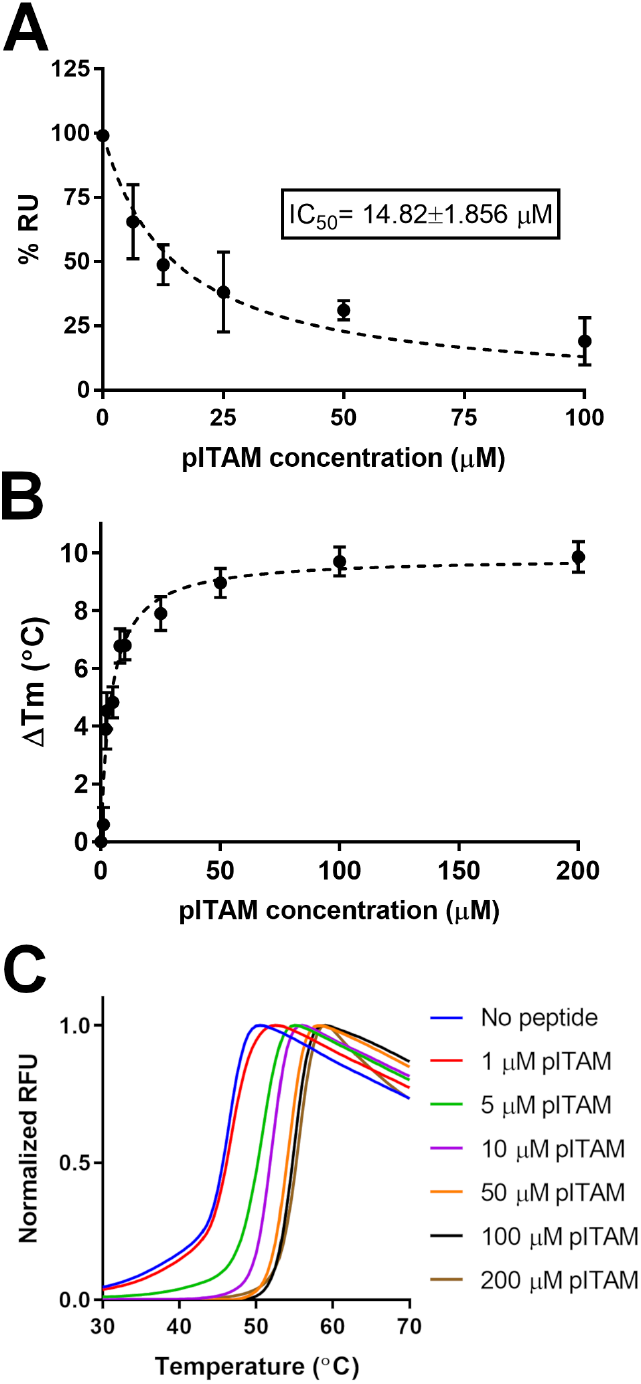
pITAM-Integrin competition experiment. **A.** Competition experiment was carried out using Surface Plasmon Resonance. The integrin β_3_ cytoplasmic domain peptide was coupled on the surface and a fixed Syk tSH2 protein concentration (25 μM) mixed with different pITAM concentrations (from 6.25 to 100 μM) was injected. The competition was evaluated considering the reduction of the Response Units (RU) at the end of the injection. The results were plotted as percentage (%) of the RU obtained in absence of pITAM peptide *versus* the pITAM concentrations (μM) used. The error bars in the graphs represent differences between three separate experiments. **B and C**. An estimation of pITAM binding to Syk tSH2 based on thermal denaturation experiment. B shows a the change of Tm in the presence of different concentrations of pITAM. C shows the actual fluorescence thermal denaturation data.

## Discussion

In this study we wanted to test the hypothesis that the integrin binding surface on the regulatory domain of Syk is different from the pITAM binding site. We were able to confirm that the main binding site on integrin β_3_ subunit cytoplasmic domain was the IA segment of Syk, but the N-terminal SH2 domain was also found to contribute to the binding. This is clearly a different binding mode than for pITAM. However, contrary to our initial hypothesis, we found that the NMR chemical shift changes on Syk N-terminal SH2 domain induced by integrin partially overlapped with those of pITAM. Furthermore, pITAM competed for β_3_ integrin binding to the regulatory domain of Syk. These three main findings are discussed below.

The role of the interdomain A (IA) segment of Syk and ZAP70 in integrin binding has been reported before (13). Our results are consistent with these results. However, the more detailed binding site of integrin on Syk has not been mapped before. We used NMR spectroscopy to map the chemical shift changes caused by soluble β_3_ peptide on the N-SH2 domain. As the affinity of the β_3_ integrin peptide to N-SH2 alone is rather low and because of limited solubility of the peptide, the experiment could not be done with the N-SH2 alone. Instead, we used the N-SH2+IA fragment that has higher affinity for integrin. Even though we were not able to assign the IA segment residues in the NMR spectra, we detected several chemical shift changes in the unassigned NMR signals, presumably arising from the IA segment. In addition to this, we saw clear chemical shift changes on the assigned part of the spectra and these changes partially overlapped with to the pITAM binding surface in the available crystal structure Syk N-SH2 (9). To verify this, we also performed NMR titrations with single-phosphorylated pITAM peptide on N-SH2 and verified the interaction site of pITAM. Particularly, it is interesting that integrin titration induced chemical shift changes at residues Ser44, Arg45, Asn46 and Tyr47 that are at edge of the pTyr binding pocket. The main positively charged side chains interacting with the phosphate group of the pITAM peptide are those of Arg22 and Arg42. Asn46 may also have a polar interaction with the phosphate and thus it is interesting to note that its side chain NHδ also shifted by integrin binding. Earlier it has been shown that mouse Syk Arg41Ala (corresponding to Arg42 in human) mutation disrupts pTyr binding, but does not affect integrin signaling in platelets (21), nor the direct interaction with β_3_ (12). This is in line with our result that the environment of Arg42 does not change upon integrin titration. We observed a change in the main chain NH chemical shift of Arg22 associated with the integrin binding, but not in its side chain N^ε^H^ε^ resonance. This suggest that the positively charged side chains of Arg22 and Arg42 do not contribute to integrin interaction. Similarly, β_3_ peptide did not cause CSPs at the hydrophobic interaction site of pITAM (Gly98 and Val100), further demonstrating the specificity of our titration.

The observed integrin-responsive surface on Syk N-SH2 is located close to the IA segment in the available structure of the Syk regulatory domains (Figure 5A). This fits with IA and N-SH2 being both involved in the interaction with a linear integrin peptide. Interestingly this surface is largely masked by the C-SH2 domain in the pITAM Syk tandem SH2 complex crystal structure (Figure 5B). On the other hand, in the structure of full-length Syk (10) this surface is mostly available in a groove between the IA segment and the N-SH2 domain (Figure 5A). This suggests that when pITAM binds to the regulatory domain of Syk, either pTyr interaction on N-SH2 or the movement of C-SH2 domain and IA segment might hinder integrin binding on N-SH2.

**Figure 5:**
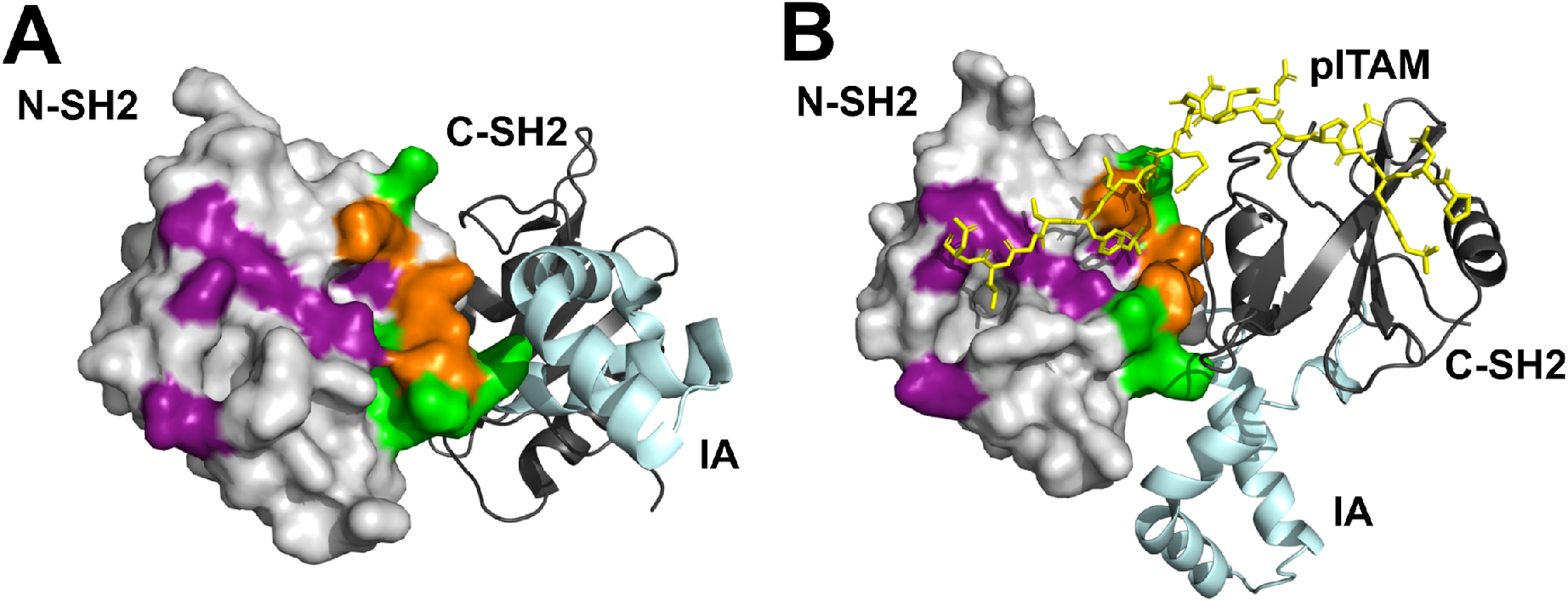
Location of the IA segment and C-SH2 in relation to the integrin-responsive surface of N-SH2. Comparison of the tSH2 structure from the full length Syk (PDB 4FL2) (**A**) and in the pITAM-tSH2 complex (PDB 1A81) (**B**). The residues whose chemical shifts changes upon both integrin on pITAM titration are coloured orange, those changed by only integrin green, and only pITAM purple. Note that in the conformation found in the full length Syk (A), the IA segment is located very close to the integrin-responsive surface, suggesting that the IA segment may be partly contributing to to the changes. On the other hand, in the presence of pITAM the surface is mostly covered by the C-SH2 and pITAM peptide (B).

In line with the NMR titration experiments, we found that soluble pITAM peptide inhibit the interaction of Syk regulatory domain with the surface coupled integrin in the SPR assay. The apparent IC50 value was 15 μM, which is much higher concentration than the measured K_D_ between pITAM and tSH2 (about 100 nM; (10, 18)). However, the measured IC50 value was close to the K_D_ value that we estimated using a thermal stabilization assay. K_D_ values of pITAM-Syk interaction vary considerably depending on the assay and proteins used, for instance Grädler et al. reported K_D_ of 100 nM by isothermal titration calorimetry and 50 μM by SPR (10). This may reflect the conformational flexibility of Syk.

In contrast to our current result, Woodside and collaborators observed no cross-inhibition between pITAM and β_3_ integrin peptides in an enzyme-linked immunosorbent assay where His6-integrin peptide was allowed to bind surface immobilized GST-Syk(6-370), or in pull-down experiments between His6-integrin coupled to beads and binding of soluble GST-Syk(6-370) (12). Although we cannot explain the cause of the different results, it is possible that differences in pITAM peptides used in the assay explain the variation. Woodside et al., used FcεRIγ pITAM at 10 μM, we used CD3ε pITAM at 1-100 μM.

In conclusion, our results suggest that Syk cannot bind to integrins and pITAM at the same time. This fits with the finding that in platelets integrin-mediated Syk signaling has been found to be independent on pITAM signaling (21). On the other hand, in neutrophils fibrinogen and β_3_ - induced oxidative burst is dependent on ITAM signaling (22). Similarly in neutrophils and macrophages, integrin-induced Syk phosphorylation also requires ITAM signaling (3). Our current results imply that in those cases where ITAM and integrin-induced activation of Syk are both required, they need to occur spatially or temporally separated.

## Experimental procedure

### Proteins expression and purification

Tandem SH2 (amino acids 9-264) and shorter constructs (Figure 1) were cloned in plasmid pGTvL1-SGC (Structural Genomincs Consortium, University of Oxford) using a ligation-independent method (23). The final construct was sequence verified. The final proteins contain N-terminal Ser-Met sequence derived from the vector. Proteins were expressed using *Escherichia coli* BL21 gold strain (Agilent Technology Inc.) cultured in Terrific Broth (TB) at 25 °C for 20 hours and induced at OD600=0.8 using 1 mM isopropyl-β-D-1-thiogalactopyranoside. Cells were lysed in cold using a French press. Proteins were captured using glutathione-agarose column (Protino glutathione-agarose 4B, Macherey-Nagel Gmbh) equilibrated with 20 mM Tris HCl, 100 mM NaCl, 1 mM DTT, pH ranging from 7 to 8 depending on the isoelectric point of the specific protein construct. Glutathione-S-Transferase-protein complex was cut using TEV protease (Invitrogen), and further purified with size-exclusion chromatography with HiLoad 26/60 Superdex 75 column (GE Healthcare). The final protein preparations were concentrated to 1 mM using Amicon^®^ Ultra 10 or 3 kDa cut off centrifugal filters (Merck KGaA). Aliquots were frozen in liquid nitrogen and stored in −80 °C.

### Surface Plasmon Resonance (SPR)

The SPR experiment was done using Biacore X instrument (GE Healthcare Inc.). Integrin β_3_ cytoplasmic tail peptide (Uni-Prot ID: P05106 CKFEEERARAKWDTANNOLYKEATSTFTNI TYRGT, ProteoGenix, Schiltgheim, France) was coupled onto CM5 chip (GE healthcare) using thiol coupling). The coupling procedure was done as previously described (20). The experiments were carried out using 10 mM HEPES, pH 7.5 as running buffer, with a flow rate of 10 μl/min using two flow cells. 60 μl of proteins dilutions were injected followed by a 150 s delayed wash. 2-fold protein dilutions were prepared from 100 to 1.6 μM, except for N-SH2 where 300, 200 and 150 μM were added due to the low affinity binding. No surface regeneration was done between the measurements. The plotted response units (RU) were obtained by subtraction of the response obtained from the non-coupled reference sensor from the signal obtained from integrin-functionalized sensor. The curves were fitted using Langmuir binding model with Biacore Evaluation software 3.1 provided by the manufacturer.

### pITAM-Integrin competition assay with SH2 constructs

pITAM competition assay was performed using SPR. Protein concentration was kept constant equal to 25 μM throughout the experiment and different concentrations of pITAM peptide from CD3ε chain (UniProt ID: P07766, NPDpYEPIRKGQRDLpYSGLNQR, ProteoGenix) ranging from 6.25 to 100 μM were added to the protein, incubated for 5 min at room temperature and, then, injected on the sensor chip. A sample without peptide was also run. The final response units (RU) were obtained by subtraction of the response obtained from the non-functionalized sensor from the response obtained from the integrin-functionalized sensor. The inhibition effect was calculated considering the reduction of the RU at the end of the injection compared to the RU obtained in the absence of pITAM. Each measurement was repeated three times, the average and the standard deviation were calculated. Data were plotted using the reduction of RU expressed as percentage of the response obtained without the pITAM *versus* the pITAM concentration used. Data were fit with GraphPad Prism 7 software using “[Inhibitor] *vs*. normalized response” model to obtain the IC50.

### Nuclear Magnetic Resonance (NMR) spectroscopy

Expression of ^15^N-labelled or ^15^N,^13^C-labelled N-SH2 or N-SH2+IA was done using *Escherichia coli* BL21 gold strain cultured in M9 medium supplemented with ^15^N-NH4Cl, ^13^C-Glucose (Cambridge Isotope Laboratories, Inc) and trace elements at 25 °C for 20 hours using 1 mM isopropyl-β-D-1-thiogalactopyranoside. The purification was done as described above. For the NMR experiments the sample was concentrated to 50 μM buffered with 50 mM Na-Phosphate pH 5 and supplemented with 4% v/v D2O. The double resonance ^15^N-HSQC was recorded at 25 °C using Bruker Avance III HD 800 MHz NMR spectrometer, equipped with cryogenically cooled TCI ^1^H, ^13^C, ^15^N triple resonance probe head. The peptide titrations were performed by adding increasing amount of peptide to the protein preparation. Integrin β_3_ peptide was titrated with N-SH2+IA protein construct using protein:peptide ratios 1:1, 1:2.5, 1:5 and 1:10. N-SH2 was used in the titration of CD3ε single pTyr ITAM peptide (NPDYEPIRKGQRDLpY SGLNQR, CASLO) with protein:peptide ratios 1:40 and 1:80.

### Thermal shift assay

The thermal shift assay was performed using thermal cycler C1000 Touch (BioRad) in 96 well plate (Hard-Shell^®^ PCR plates, BioRad). 4 μM of all studied protein constructs were mixed with different concentrations of peptides ranging from 1 to 200 μM. Sypro Orange protein gel staining 5000x concentrated (Invitrogen) was used as fluorescent dye at the concentration suggested from the manufacturer; the buffer used in the assay mix was 10 Mm Hepes, 20 Mm NaCl, 10 mM MgCl_2_, 1 mM EGTA, pH 7.4. The final volume of the mixture was 25 μl. A blank containing the buffer was run as well as a sample containing the protein without the peptides and one with the peptides alone. The temperature was raised from 20 to 95 °C with steps of 0.5 °C and the temperature was kept constant for 30 seconds before recording the fluorescence signal. Each measurement was run in quadruplicate. The final fluorescence curve was obtained subtracting the signal obtained from the buffer in the case of the proteins without the peptide and subtracting the buffer and the peptides signal from the sample where the proteins were mixed with the peptides. The fluorescence curves were then fitted with Boltzmann isotherm (24) using GraphPad Prism 7 to calculate the melting temperature (Tm). Tm was plotted *versus* the pITAM concentrations used and the data were fitted using models “One-site-specific binding” and “Two site-specific binding” with GraphPad Prism 7 to calculate the dissociation constant K_D_.

## Acknowledgements

The authors acknowledge Mr Petri Papponen for technical help. This work has been supported by Academy of Finland Grants 278668 (to J.Y.), 290506 (to V.P.H), and 288235 (to P.P).

---

**Supporting Information figure.**
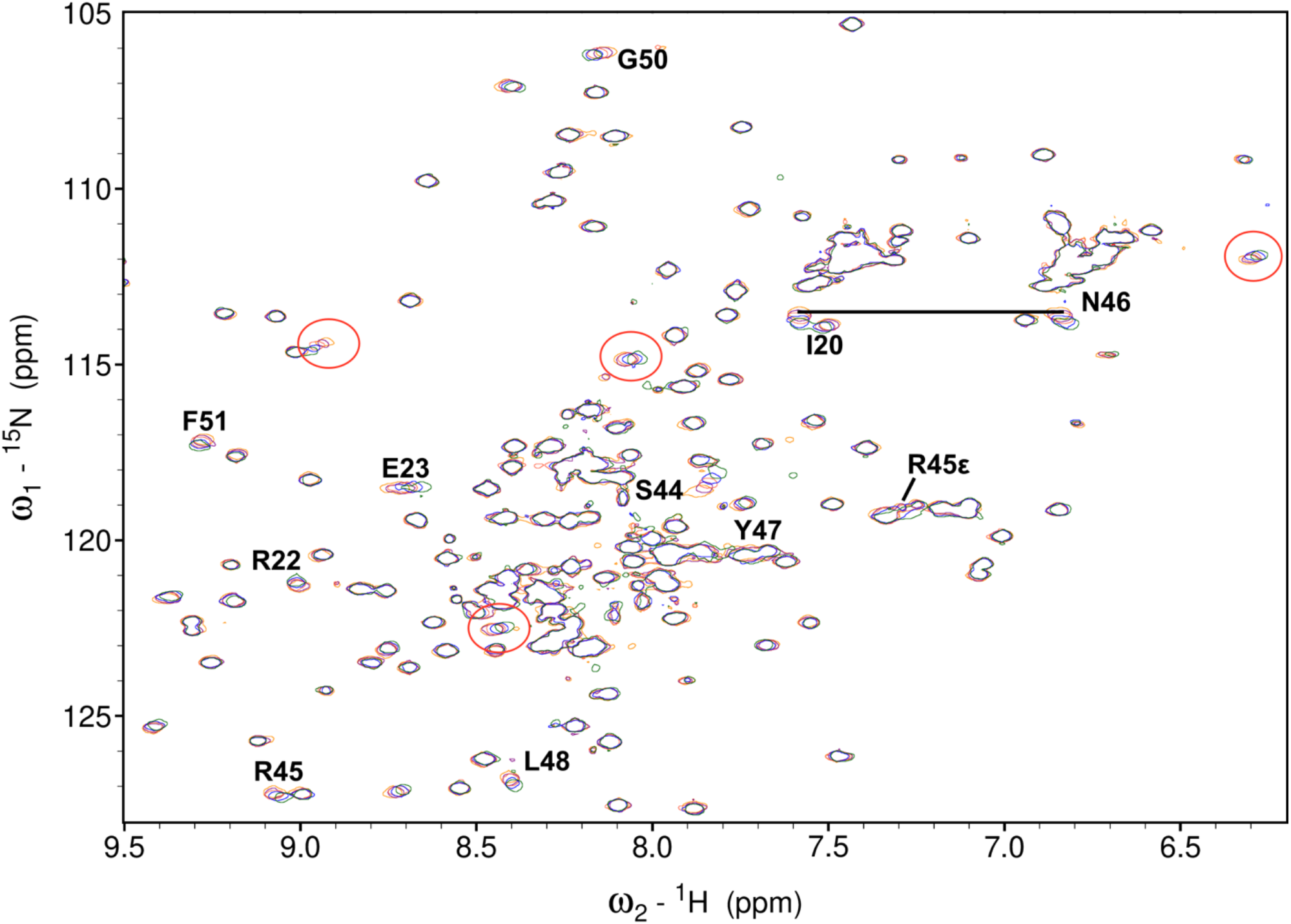
Integrin titration NMR spectra. A region of the ^15^N-HSQC spectrum of Syk N-SH2+IA fragment without integrin peptide (red) and in the presence of 1x (orange), 2.5x (yellow) and 10x (blue) excess of the integrin peptide. The x-axis corresponds to proton chemical shift, and and the y-axis to ^15^N chemical shift. The peaks indicated with red circles are unassigned peaks that change upon titration and apparently derived from the IA segment. The assigned peaks from N-SH2 that show most prominent changes are labelled. The two peaks from the Asn46 side chain are connected with a horizontal line.

